## Supplementary Information for "Effective CRISPRa-Mediated Control of Gene Expression in Bacteria Must Overcome Strict Target Site Requirements"

**This PDF file includes:**

- Supplementary Figures 1-9
- Supplementary Tables 1-5
- Supplementary Methods, including reporter gene sequences
- Supplementary References

### Supplementary figures

### Figure S1

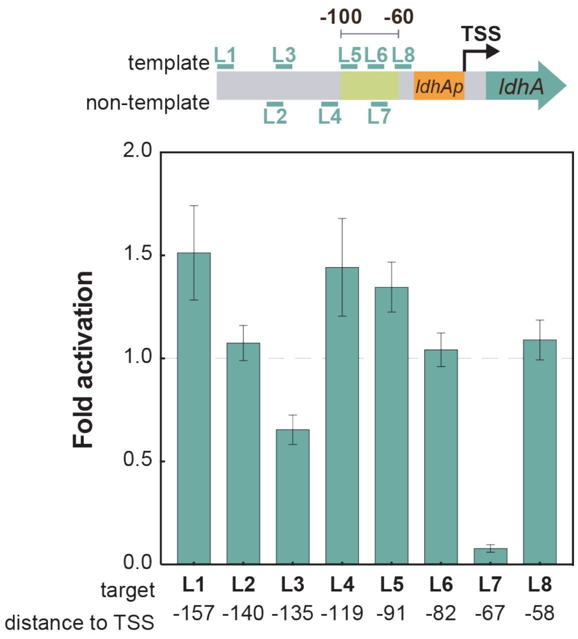


**Supplementary Figure 1: CRISPRa at the endogenous gene target *ldhA* does not follow predicted trends.**

Eight scRNA target sites (L1-L8) upstream of the *ldhA* promoter were selected. Three of the target sites (L5-L7) were within the 40 bp window where CRISPRa is effective (-100 to -60). While L1, L4, and L5 resulted in weak increases in gene expression, there was no apparent relationship between the position of the sites and *ldhA* expression levels. Gene expression was measured using RT-qPCR. Fold activation represents expression levels relative to an off-target control (hAAVS1). Bars indicate the average values between three technical replicates and black dots indicate the values of individual replicates. Error bars indicate the standard error of the mean between three technical replicates.

### Figure S2

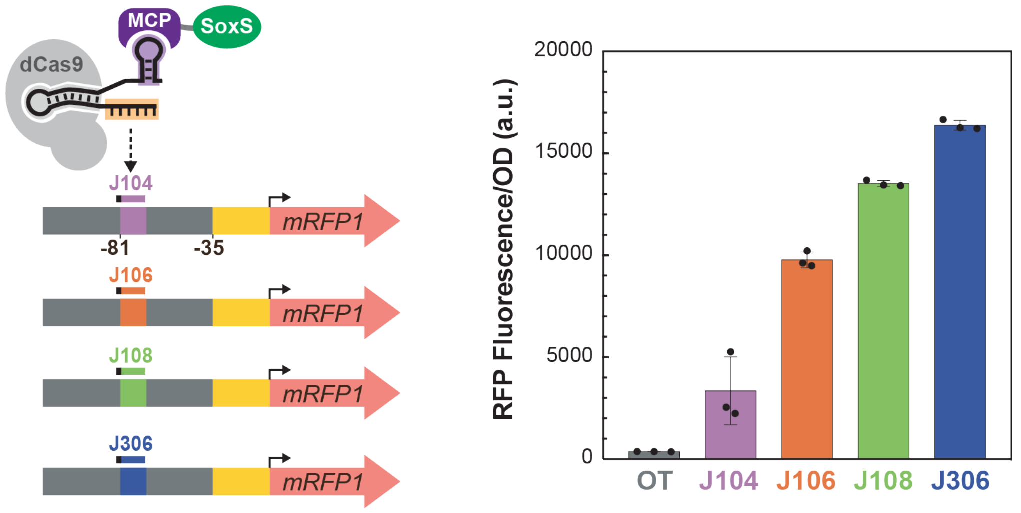


**Supplementary Figure 2: CRISPRa activity depends on the target sequence on the scRNA.** Reporter cassettes that differ only by the sequence of the 20 base scRNA target site give a broad range of gene expression levels, demonstrating that the sequence of the scRNA target site can have a substantial effect on CRISPRa. Three new reporter plasmids were constructed where the J306 target site, located at -81 from the TSS, on the J3-J23117-mRFP1 reporter was replaced by the J104, J106, and J108 sequence. Activation at each promoter was tested when CRISPRa was targeted to their cognate scRNA site. The off-target negative control (OT) represents a strain expressing the original reporter with the J306 site and the CRISPRa components to target an off-target site (J206). Fluorescence/$OD_{600}$ values were measured using a plate reader. Bars indicate the average values between three biological replicates. Black dots indicate the values of individual biological replicates. Error bars indicate the standard deviation between biological replicates.

### Figure S3

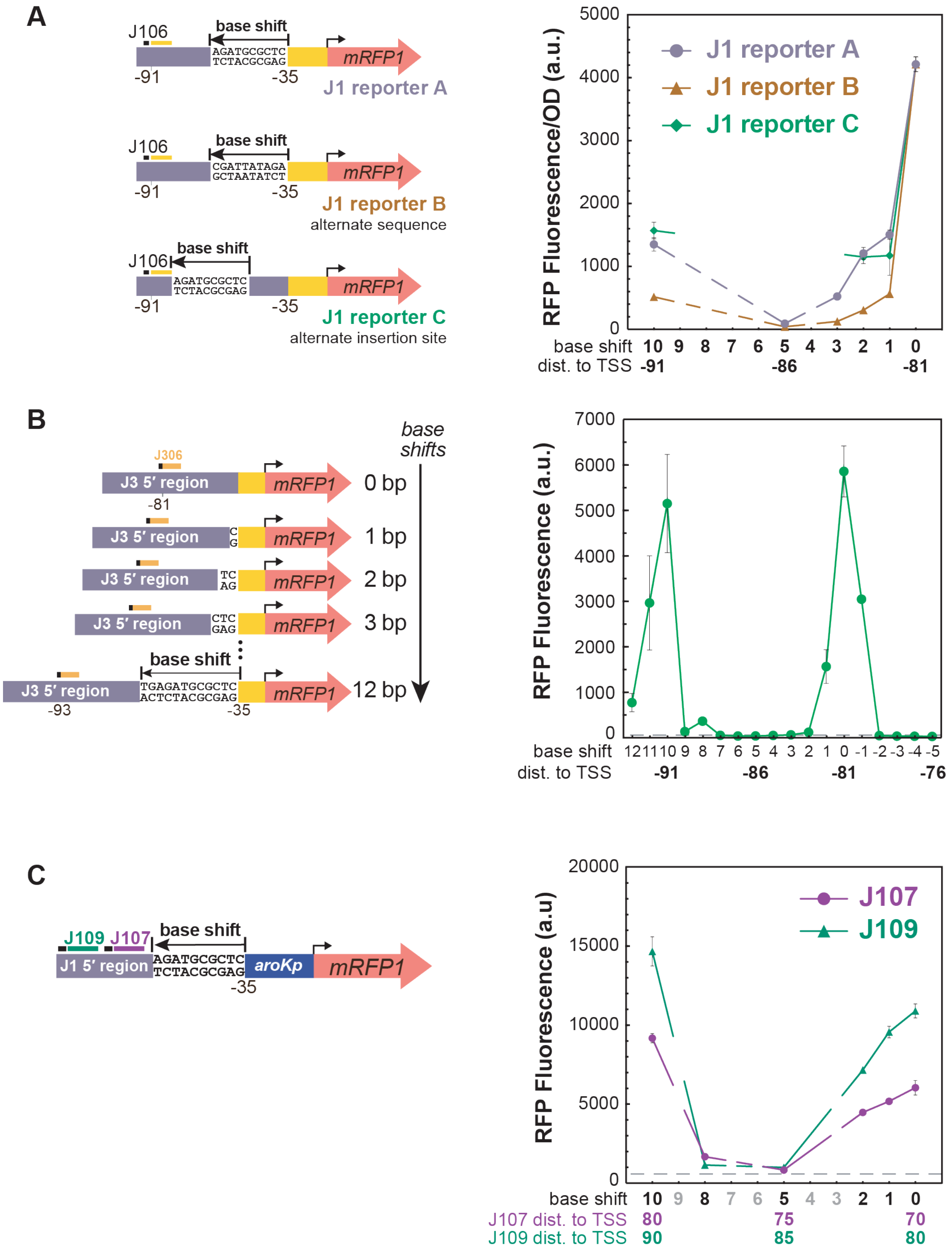

A
J106
base shift
AGATGCGCTC
TCTACGCGAG
mRFP1
-91
-35
J1 reporter A
J106
base shift
CGATTATAGA
GCTAATATCT
mRFP1
-91
-35
J1 reporter B
alternate sequence
base shift
J106
AGATGCGCTC
TCTACGCGAG
mRFP1
-91
-35
J1 reporter C
alternate insertion site
RFP Fluorescence/OD (a.u.)
J1 reporter A
J1 reporter B
J1 reporter C
5000
4000
3000
2000
1000
0
base shift 10 9 8 7 6 5 4 3 2 1 0
dist. to TSS -91 -86 -81
B
base shifts
J306
J3 5′ region
mRFP1
0 bp
-81
J3 5′ region
mRFP1
1 bp
J3 5′ region
mRFP1
2 bp
J3 5′ region
mRFP1
3 bp
base shift
J3 5′ region
TGAGATGCGCTC
ACTCTACGCGAG
mRFP1
12 bp
-93
-35
RFP Fluorescence (a.u.)
7000
6000
5000
4000
3000
2000
1000
0
base shift 12 11 10 9 8 7 6 5 4 3 2 1 0 -1 -2 -3 -4 -5
dist. to TSS -91 -86 -81 -76
C
J109 J107
base shift
J1 5′ region
AGATGCGCTC
TCTACGCGAG
aroKp
mRFP1
-35
RFP Fluorescence (a.u)
J107
J109
20000
15000
10000
5000
0
base shift 10 9 8 7 6 5 4 3 2 1 0
J107 dist. to TSS 80 75 70
J109 dist. to TSS 90 85 80

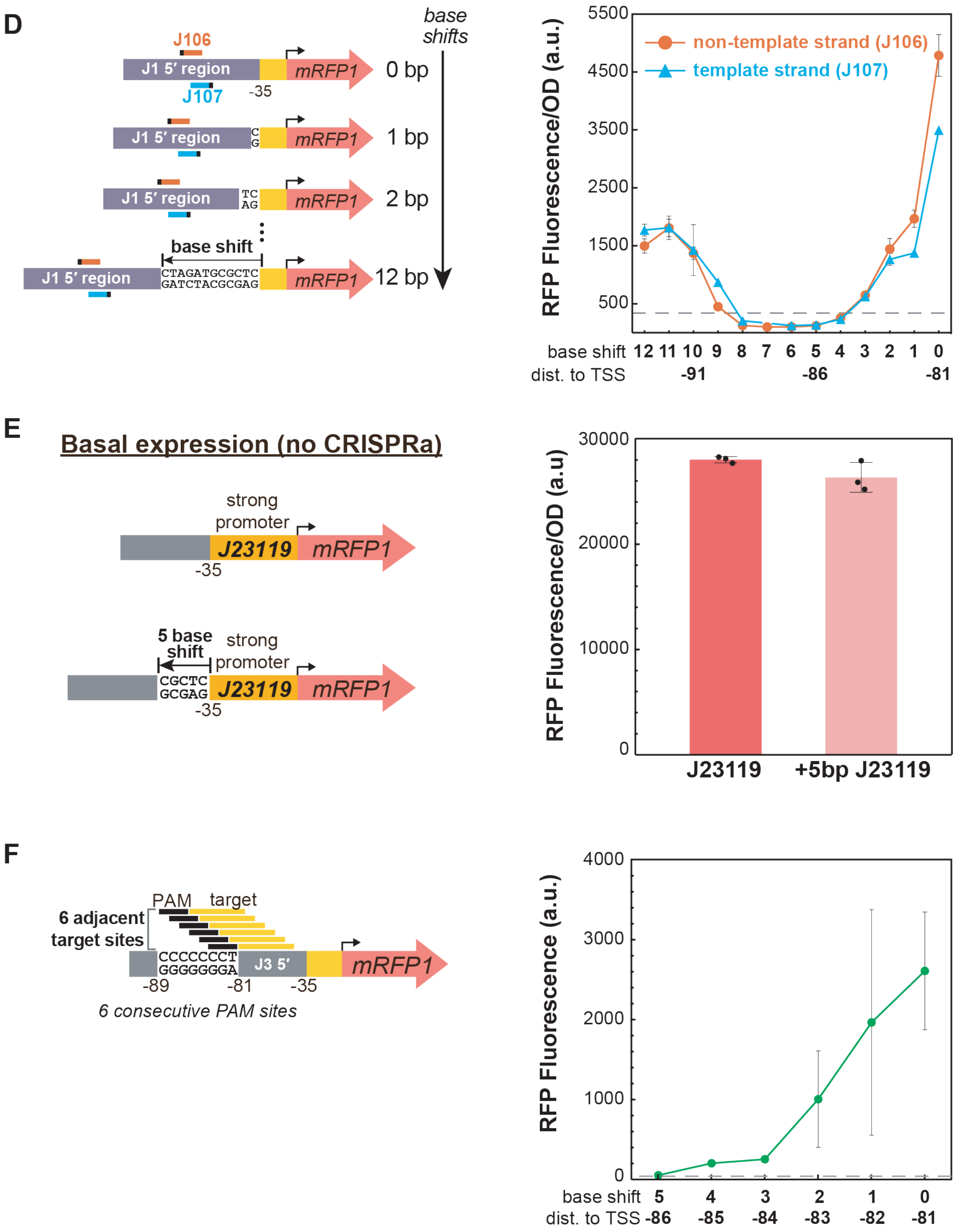


**Supplementary Figure 3: The sharp positioning dependence of CRISPRa is observed across multiple promoters.**

**A)** The sharp positioning requirements of CRISPRa are not significantly affected by the location or composition of the inserted sequence. Reporters were based on the J1-J23117-mRFP1 (Supplementary Figure 3A) with base shifts introduced in different ways. In the J1 reporter A, bases were inserted upstream of the -35 region. In the J1 reporter B, a different sequence was inserted at the same site. In the J1 reporter C, bases were inserted downstream of the J106 target site. There were modest differences between reporter A and reporter B that could indicate a contribution from sequence composition. **B)** The sharp positioning requirements of CRISPRa were observed with a different heterologous promoter. CRISPRa was targeted at the -81 site on the J3-J23117 promoter that has a different upstream sequence and a different scRNA target site (J306) compared to J1-J23117. Peaks in gene expression were observed at -81 and -91 to the TSS, displaying a 10 bp periodicity. The peaks of gene expression on the J3-J23117 promoter were sharper than in the J1-J23117 promoter (Figure 4A); gene expression decreased to baseline levels after shifting only 2 bp from the peak position. After a complete 10 bp-shift period, CRISPRa activity was fully restored to the original peak expression. Reporter gene sets were constructed by inserting 0-12 bp or deleting 1-5 bp upstream of the -35 of the J3-J23117-mRFP1 reporter. The grey line represents the baseline activity of the J3-J23117-mRFP1 reporter strain containing an empty vector instead of the CRISPRa component plasmid. For comparison, previous CRISPRa data at -81 and -91 are shown on the schematic above the plot[1]. **C)** The sharp positioning requirements of CRISPRa are observed when targeting a different minimal promoter. The J1-*aroKp2* promoter displayed positioning requirements similar to the J1-J23117 promoter. Decreases in CRISPRa activity after 3 bp shifts and recovery at a 10 bp shift were observed on both the J107 and the J109 target sites. The J1-*aroKp2* promoter was constructed by replacing the BBa_J23117 minimal promoter from the J1-J23117 promoter to the aroKp2 minimal promoter. A J1-*aroKp2*-mRFP1 reporter series was constructed by adding 0 bp, 1 bp, 2 bp, 5 bp, 8 bp, and 10 bp upstream of the -35 region. **D)** The sharp positioning requirements of CRISPRa are observed on both the template and the non-template strand. For both the J106 target site on the non-template strand and the J107 target on the template strand, the CRISPRa activity decreased to baseline levels after shifting 3 bp from its original position and recovered after shifting 10 bp. The reporter plasmid was the same as in Figure 4A. The dotted grey line indicates the negative control where an off-target scRNA (J206) was co-transformed with the original reporter with no inserted bases. **E)** Adding 5 bases adjacent to the -35 region does not dramatically alter the expression of the promoter. The 5 bases added were the same as those used to shift the J1 and J3 promoters (Figure 4A and S3A). This experiment was performed using a strong minimal promoter (BBa_J23119), so that any detrimental effects would be detectable. **F)** The sharp positioning requirements of CRISPRa were observed when tested in a single reporter with multiple consecutive PAM sites. 6 consecutive PAM sites on the non-template strand were introduced by placing a CCCCCCCT sequence between -89 and -81 bp to the TSS on the J3-J23117-mRFP1 reporter. Maximum gene expression was observed at the original -81 site, after which expression gradually decreased to one third of the maximum activity after moving 2 bp away (-83). Gene expression decreased further when the scRNA target was moved 3 bp and 4 bp away from the TSS (-84 and -85), and reached the baseline when moved 5 bp (-86). The grey line represents the baseline activity of the reporter strain containing an empty vector instead of the CRISPRa component plasmid. Fluorescence/$OD_{600}$ values in panels A, D, and E were measured using a plate reader. Median fluorescence values in panels B, C, and F were measured using a flow cytometer. Colored dots and bars indicate the average values between three biological replicates. Black dots indicate the values of individual biological replicates. Error bars indicate the standard deviation between biological replicates.

### Figure S4

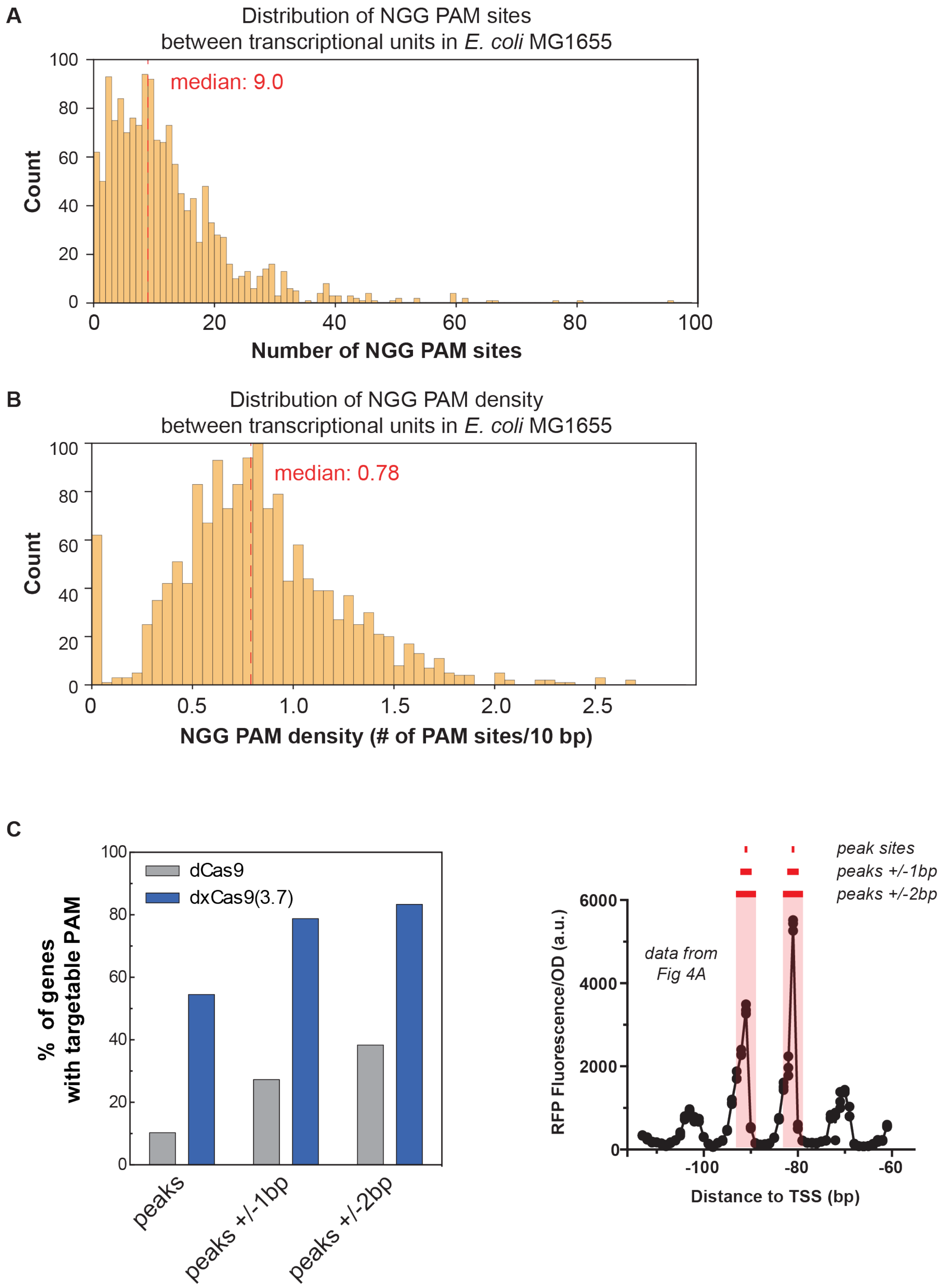


**Supplementary Figure 4: Availability of PAM sites between transcriptional units in *E. coli* MG1655.**

**A)** Distribution of the number of NGG PAM sites between transcriptional units in *E. coli* MG1655. The median number of NGG PAM sites between transcriptional units in *E. coli* MG1655 is 9.0. Methods for extracting the sequence of the DNA regions between transcriptional units in *E. coli* MG1655 and identifying available PAM sites are described in the Supplementary Methods. **B)** Distribution of the density of NGG PAM sites between transcriptional units in *E. coli* MG1655. The density of PAM sites is reported over a 10 bp window. This window size was chosen because it corresponds to a full turn of the DNA helix. The density of PAM sites in a 10 bp window was calculated for each sequence between transcriptional units as the total number of PAM sites in the sequence divided by the length of the intergenic sequence and then multiplied by 10. The median density of NGG PAM sites per 10 bp between transcriptional units in *E. coli* MG1655 is 0.78. **C)** The number of genes in the *E.coli* genome that can be targeted by dCas9 is small, and the expanded PAM variant dxCas9(3.7) significantly increases the number of genes with predicted effective target sites for CRISPRa. The plot to the left shows percentage of genes with at least one PAM site targetable by dCas9 or dxCas9(3.7) at: the positions where CRISPRa displays a peak in activity. The plot to the right illustrates the range of positions on the non-template strand chosen for the analysis. Corresponding peaks on the template strand were also included. Analyses were performed using data generated when selecting candidate endogenous genes for activation (Supplementary Methods).

### Figure S5

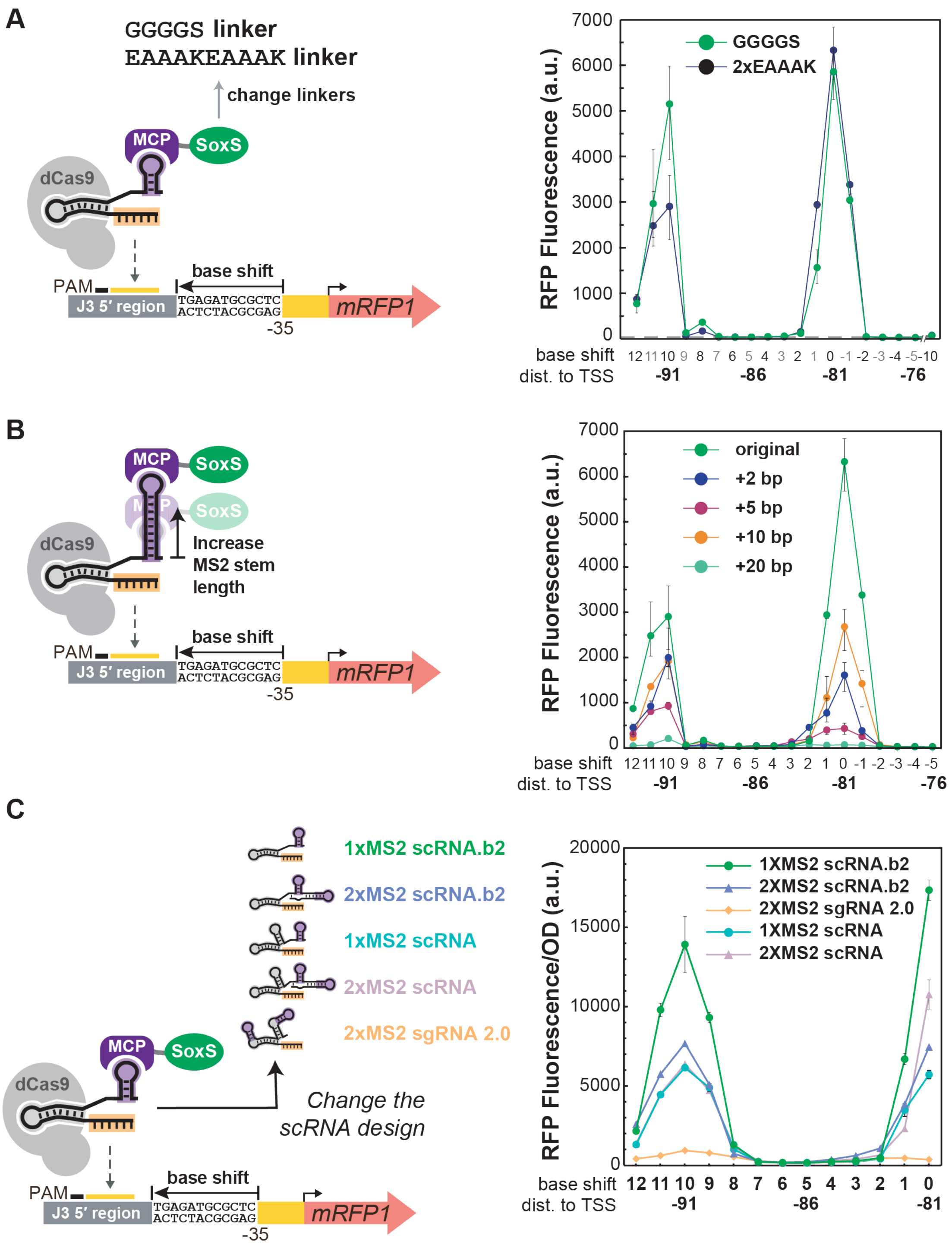


**Supplementary Figure 5: Modifying the CRISPRa complex structure does relax the sharp positioning requirements of CRISPRa.**

**A)** Changing the linker between MCP and SoxS does not change the positioning dependence of CRISPRa. A CRISPRa complex with the MCP-(EAAAKEAAAK)-SoxS(R93A/S101A) activation domain displayed 10 bp periodicity of peak expression similar to that with the MCP-(GGGGS)-SoxS(R93A/S101A) activation domain. The EAAAKEAAAK linker is predicted to be more rigid than the GGGGS linker[2]. The grey line represents the baseline activity of the J3-J23117-mRFP1 reporter strain containing an empty vector instead of the CRISPRa component plasmid. **B)** Extending the length of the MS2 stem does not change the positioning dependence of CRISPRa. CRISPRa systems with scRNAs that have +2, +5, and +10 RNA base pairs added to the bottom of the MS2 stem displayed the same 10 bp periodicity but lower peak activity compared to the original 1xMS2 scRNA.b2. The strain having the scRNA with +20 bp extended MS2 stem did not show any CRISPRa activity. **C)** No improvements in the effective target range were observed for CRISPRa systems with alternative scRNA designs. The scRNA designs tested are: 1xMS2 scRNA.b2 and 2XMS2 scRNA.b2 with the tracrRNA hairpin removed[1], the original 1xMS2 and 2XMS2 scRNA design having the tracrRNA hairpin[3], and sgRNA 2.0 where the MS2 hairpins are extended from the RNA stems on the sgRNA[4]. CRISPRa systems expressing a 2XMS2 scRNA.b2, 1xMS2 scRNA, and 2XMS2 scRNA displayed the same 10 bp periodicity but lower peak activity than 1xMS2 scRNA.b2. Expressing a sgRNA 2.0 resulted in no CRISPRa activity. Median fluorescence values in panels A and B were measured using a flow cytometer. Fluorescence/$OD_{600}$ values in panels C were measured using a plate reader. Dots indicate the average values between three biological replicates. Error bars indicate the standard deviation between biological replicates.

### Figure S6

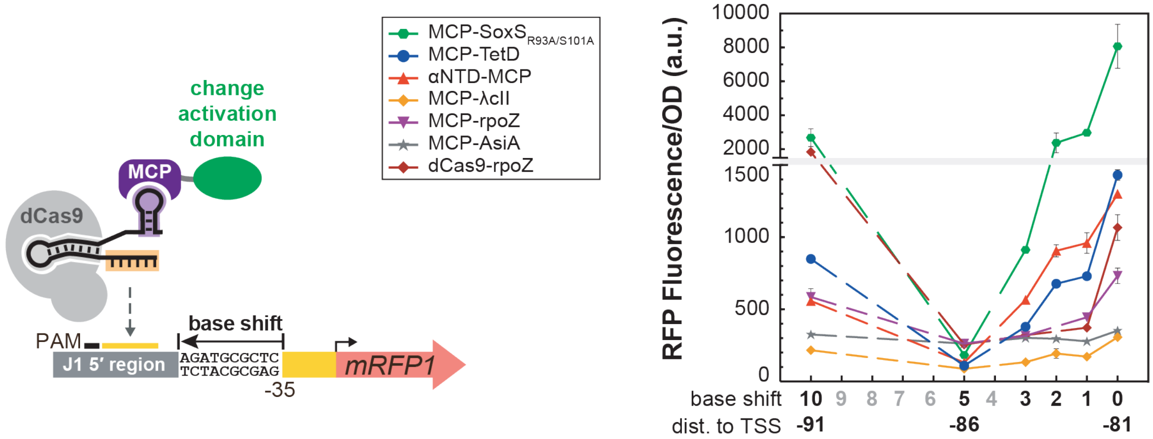


**Supplementary Figure 6: Performing CRISPRa with alternative activation domains does not expand the range of targetable positions.**

CRISPRa with MCP-TetD and αNTD-MCP activation domains[1] displayed similar positioning dependence as MCP-SoxS(R93A/S101A), where gene expression decreases as the position shifts from 0 to 3 bp, reaches the baseline at a 5 bp shift, and increases again at a 10 bp shift. CRISPRa with MCP-rpoZ[1] and dCas9-rpoZ[5] activation domains displayed more stringent positioning dependence, where gene expression approaches the baseline at 1 bp shift. All alternative activation domains gave weaker peak gene expression at 0 bp shift compared to MCP-SoxS(R93A/S101A). MCP-λcII and MCP-AsiA[1] activation domains did not show any significant CRISPRa activity. All activation domains were cloned into the CRISPRa component plasmid containing *Sp*-dCas9 and 1xMS2 scRNA.b2 targeting J106. To reduce the toxicity of AsiA, the MCP-AsiA plasmid also contains a co-expressed *rpoD(F563Y)* gene[1]. The MCP-SoxS(R93A/S101A), MCP-TetD, αNTD-MCP, MCP-λcII and MCP-AsiA plasmids were tested in *E. coli* MG1655. The MCP-*rpoZ* and dCas9-*rpoZ* plasmids were tested in the *E. coli* strain CD03 (MG1655/Δ*rpoZ*) (Supplementary Table 1)[1]. Fluorescence/$OD_{600}$ values were measured using a plate reader. Dots indicate the average values between three biological replicates. Error bars indicate the standard deviation between biological replicates.

### Figure S7

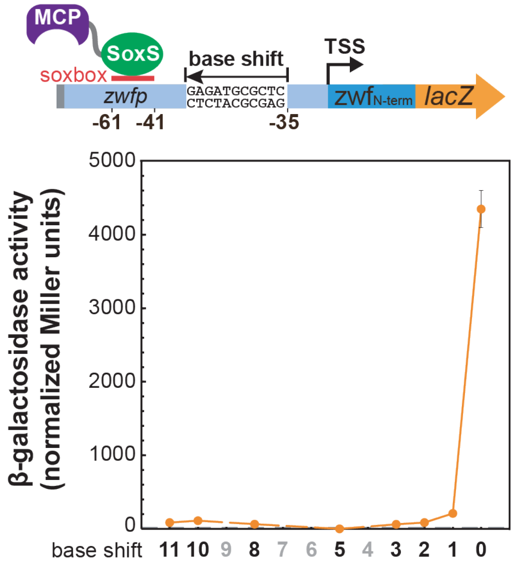


**Supplementary Figure 7: Wild type SoxS displays a sharp positioning requirement when targeting a SoxS-dependent promoter.**

When the wild type SoxS binding site (soxbox) on the endogenous *zwfp* promoter is shifted by 1 bp, gene expression decreases significantly, consistent with previous reports[6]. Shifting the soxbox further upstream causes gene expression to completely reduce to the baseline. Reporter plasmids were constructed by adding 1, 2, 3, 5, 8, 10, and 11 bp upstream of the -35 on the *zwfp-lacZ* reporter (Figure 1). Dots indicate the average β-galactosidase activity in normalized Miller units among three biological replicates and the error bars indicate the standard deviation.

### Figure S8

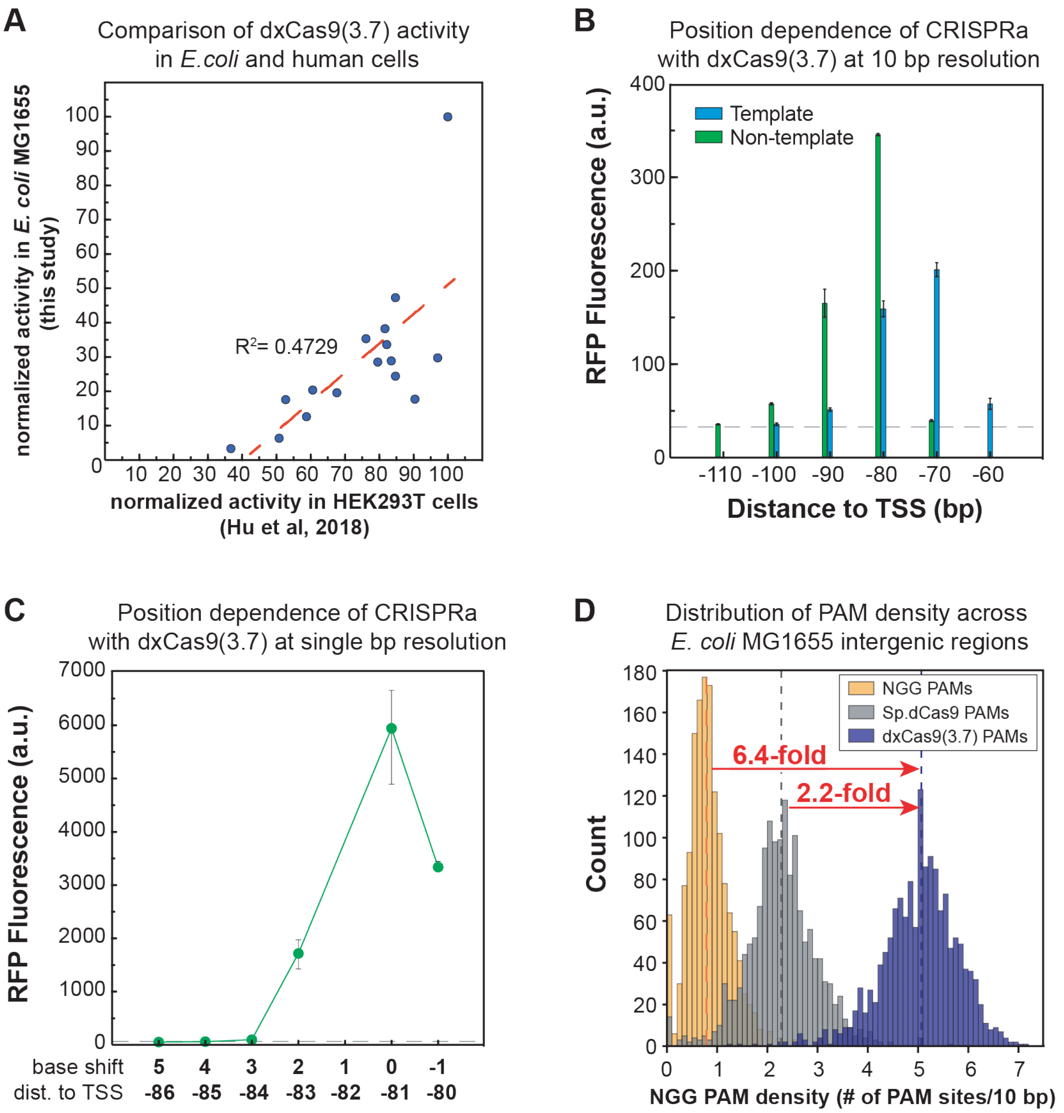


**Supplementary Figure 8: dxCas9(3.7) can target an expanded range of PAM sites and is sensitive to target site position for CRISPRa.**

**A)** Relative CRISPRa activities of dxCas9(3.7) on different PAM targets in *E. coli* correlates well with the corresponding data obtained by Hu et al.[7] in human cells. The CRISPRa activity on the NGG PAM site in both studies were normalized to 100 and the CRISPRa activity on all the other PAM sites were normalized to the value of NGG PAM site. Normalized data in this study (y axis) were plotted against the normalized data obtained by Hu et al. (x axis). **B)** CRISPRa with dxCas9(3.7) displayed similar sensitivity to target site position as *Sp*-dCas9. Peaks of activation were observed at -81 and -91 on the non-template strand and -70 and -80 on the template strand. CRISPRa was targeted to a J1-J23117-mRFP1 reporter integrated into the genome (*E. coli* CD13, Supplementary Table 1). The grey dotted line represents the baseline fluorescence of a strain containing the dxCas9(3.7) and MCP-SoxS(R93A/S101A) and an empty vector with no scRNAs. **C)** CRISPRa with dxCas9(3.7) displayed similar positioning dependence with single base shifts. Gene expression was significantly reduced when the scRNA target site was shifted 1-2 bp in either directions from the optimal position -81 bp to the TSS on the non-template strand. Shifting the scRNA target site 3-5 bp causes gene expression to fall to the baseline level. Reporter gene sets were constructed with 1 bp deleted or 2-5 bp inserted upstream of the -35 of the J3-J23117-mRFP1 reporter. The grey line represents the baseline activity of a strain containing the J3-J23117-mRFP1 reporter plasmid and a CRISPRa component plasmid with an off-target scRNA (hAAVS1). Median fluorescence values in panels B and C were measured using a flow cytometer. Bars and dots indicate the average values between three biological replicates. Error bars indicate the standard deviation between biological replicates. **D)** dxCas9(3.7) increases the probability of finding targetable PAM sites for CRISPRa. Histograms showing the distribution of the PAM density (defined in Supplementary Figure 4) between transcriptional units in *E. coli* MG1655 suggest that the average likelihood of finding a dxCas9(3.7)-compatible PAM (blue) is ~6.4-times higher than finding an NGG PAM (yellow), and ~2.2-times higher than finding a *Sp*-dCas9-compatible PAM (grey). Vertical dotted lines indicate the median PAM density for each group. The median PAM density is 5.08 for dxCas9(3.7)-compatible PAMs (NGG, AGA, AGC, AGT, CGA, CGC, CGT, GGA, GGC, GGT, TGA, TGC, TGT, GAA, GAT, CAA), 2.33 for *Sp*-dCas9-compatible PAMs (NGG, AGA, CGA, GGA, GGC, GGT, TGA). and 0.78 for NGG PAMs. Methods for extracting the sequence of *E. coli* MG1655 intergenic regions, counting targetable PAM sites and calculating the PAM density are described in the Supplementary Methods.

### Figure S9

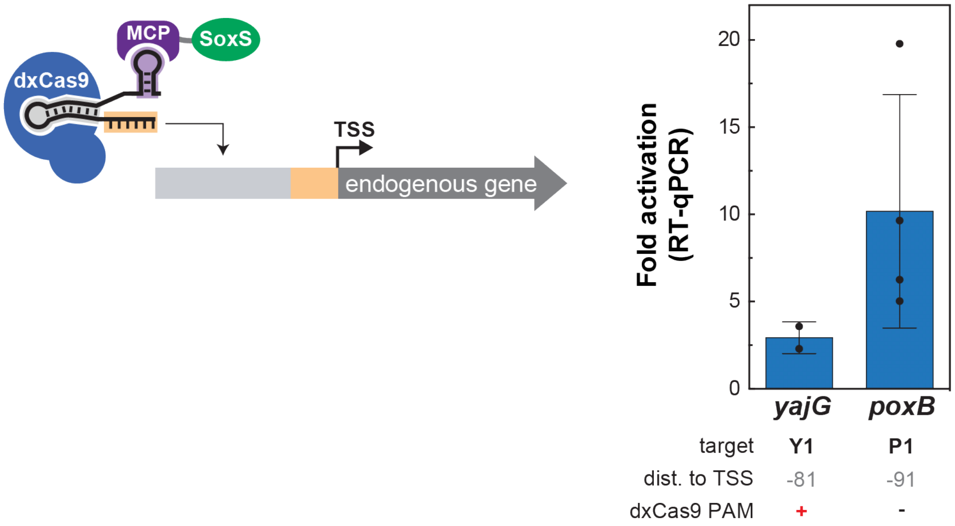


**Supplementary Figure 9. Predictive rules for CRISPRa enable activation of endogenous genes *yajG* and *poxB*.**

The scRNA sites displaying the highest activity on the *yajG* and *poxB* reporters from the *E. coli* promoter collection[8] (Figure 6B) were tested for activity with dxCas9(3.7) at the endogenous *yajG* and *poxB* genes using RT-qPCR. Fold activation represents expression levels relative to a control expressing an off-target scRNA (J306). Bars indicate the average values between two biological replicates for *yajG* and four biological replicates for *poxB* and black dots indicate the values of individual replicates. Error bars indicate the standard error of the mean.

**Supplementary tables**

**Supplementary Table 1.** *E. coli* Strains

| **Strain** | **Description** | **Genotype** | **Reference** |
|---|---|---|---|
| MG1655 | parent *E. coli* strain | F- λ- ilvG- rfb-50 rph-1 | |
| CD03 | MG1655 with *rpoZ* knocked out | MG1655 Δ*rpoZ* | 1 |
| CD06 | MG1655/sfGFP (weak promoter) | MG1655 *W1-BBa_J23117-sfGFP KanR::nfsA* | 1 |
| CD13 | MG1655/mRFP1 (weak promoter) | MG1655 *J1-BBa_J23117-mRFP1::nfsA* | This study, Suppl. Fig. 8 |

**Supplementary Table 2.** Description of the CRISPRa systems used in each figure

| Figure | Cas protein | scRNA design | scRNA target | Activation domain |
|---|---|---|---|---|
| 1B | dCas9 | 1xMS2 scRNA.b1 | W108 | MCP-(5aa)-SoxS(wild-type or mutant) |
| 2A | dCas9 | 1xMS2 scRNA.b1 | A1-A4, hAAVS1 | MCP-(5aa)-SoxS(R93A/S101A) |
| 2B | dCas9 | 1xMS2 scRNA.b2 | C1-C5, hAAVS1 | MCP-(5aa)-SoxS(R93A) |
| 3A | dCas9 | 1xMS2 scRNA.b2 | J306, J206 | MCP-(5aa)-SoxS(R93A) |
| 3B | dCas9 | 1xMS2 scRNA.b2 | J101-J120, hAAVS1 | MCP-(5aa)-SoxS(R93A/S101A) |
| 3C | dCas9 | 1xMS2 scRNA.b2 | J306, J206 | MCP-(5aa)-SoxS(R93A) |
| 3D | dCas9 | 1xMS2 scRNA.b2 | J306, J206 | MCP-(5aa)-SoxS(R93A/S101A) |
| 4A | dCas9 | 1xMS2 scRNA.b2 | J102, J104, J106, J108, J110, J206 | MCP-(5aa)-SoxS(R93A) |
| 4B | dCas9 | 1xMS2 scRNA.b2 | J106 | MCP-(5aa)-SoxS(R93A), MCP-(10aa)-SoxS(R93A), MCP-(20aa)-SoxS(R93A) |
| 5A | dCas9 / dxCas9(3.7) | 1xMS2 scRNA.b2 | J306 | MCP-(5aa)-SoxS(R93A/S101A) |
| 5B | dCas9 / dxCas9(3.7) | 1xMS2 scRNA.b2 | M1, M2, J206 | MCP-(5aa)-SoxS(R93A/S101A) |
| 6A | dCas9 / dxCas9(3.7) | 1xMS2 scRNA.b2 | Y1-Y3, J306 | MCP-(5aa)-SoxS(R93A/S101A), tet-inducible |
| 6B | dxCas9(3.7) | 1xMS2 scRNA.b2 | Y1, Y2, P1, P2, U1, U2, D1, D2, B1, B2, E1, E2, J306 | MCP-(5aa)-SoxS(R93A/S101A), tet-inducible |

| | | | | |
|---|---|---|---|---|
| S1 | dCas9 | 1xMS2 scRNA.b2 | L1-L8, hAAVS1 | MCP-(5aa)-SoxS(R93A) |
| S2 | dCas9 | 1xMS2 scRNA.b2 | J104, J106, J108, J306, J206 | MCP-(5aa)-SoxS(R93A) |
| S3A | dCas9 | 1xMS2 scRNA.b2 | J106 | MCP-(5aa)-SoxS(R93A) |
| S3B | dCas9 | 1xMS2 scRNA.b2 | J306 | MCP-(5aa)-SoxS(R93A/S101A) |
| S3C | dCas9 | 1xMS2 scRNA.b2 | J107, J109 | MCP-(5aa)-SoxS(R93A/S101A) |
| S3D | dCas9 | 1xMS2 scRNA.b2 | J106, J107, J206 | MCP-(5aa)-SoxS(R93A) |
| S3F | dCas9 | 1xMS2 scRNA.b2 | J306, J306+1-5 | MCP-(5aa)-SoxS(R93A/S101A) |
| S5A | dCas9 | 1xMS2 scRNA.b2 | J306 | MCP-(5aa)-SoxS(R93A/S101A), MCP-(2xEAAAK)-SoxS(R93A/S101A) |
| S5B | dCas9 | 1xMS2 scRNA.b2, 1xMS2 scRNA.b2 + 2/5/10 bp MS2 stem extension | J306 | MCP-(2xEAAAK)-SoxS(R93A/S101A) |
| S5C | dCas9 | 1xMS2 scRNA.b2, 2xMS2 scRNA.b2, 2xMS2 sgRNA2.0, 1xMS2 scRNA, 2xMS2 scRNA | J306 | MCP-(5aa)-SoxS(R93A) |

| | | | | |
|---|---|---|---|---|
| S6 | dCas9 | 1xMS2 scRNA.b2 | J106 | MCP-(5aa)-SoxS(R93A/S101A), Alternative activators |
| S8B | dxCas9(3.7) | 1xMS2 scRNA.b2 | J104-J113 | MCP-(5aa)-SoxS(R93A) |
| S8C | dxCas9(3.7) | 1xMS2 scRNA.b2 | J306 | MCP-(5aa)-SoxS(R93A) |
| S9 | dxCas9(3.7) | 1xMS2 scRNA.b2 | Y1, P1, J306 | MCP-(5aa)-SoxS(R93A/S101A), tet-inducible |

**Supplementary Table 3.** gRNA Target Sites

| sgRNA target | DNA Sequence | Target Strand[a] | Distance to TSS[b] |
|---|---|---|---|
| W108 | GAAGATCCGGCCTGCAGCCA | NT | 91 |
| J101[c] | TGGGTTCCACCGGATACCTC | T | 40 |
| J103[c] | AGGCGTCCTTTGGGTTCCAC | T | 50 |
| J105[c] | CGGTTACCAAAGGCGTCCTT | T | 60 |
| J107[c] | CGGTGTCCTGCGGTTACCAA | T | 70 |
| J109[c] | AGGTATCCTGCGGTGTCCTG | T | 80 |
| J111[c] | GGGCGACCTCAGGTATCCTG | T | 90 |
| J113[c] | GGGCCACCACGGGCGACCTC | T | 100 |
| J115[c] | TGGTGACCATGGGCCACCAC | T | 110 |
| J117[c] | GGGTGACCTATGGTGACCAT | T | 120 |
| J119[c] | TGGTTGCCAAGGGTGACCTA | T | 130 |
| J121[c] | AGGACACCTTTGGTTGCCAA | T | 140 |
| J102[c] | AGGTATCCGGTGGAACCCAA | NT | 61 |
| J104[c] | TGGAACCCAAAGGACGCCTT | NT | 71 |
| J106[c] | AGGACGCCTTTGGTAACCGC | NT | 81 |
| J108[c] | TGGTAACCGCAGGACACCGC | NT | 91 |
| J110[c] | AGGACACCGCAGGATACCTG | NT | 101 |
| J112[c] | AGGATACCTGAGGTCGCCCG | NT | 111 |
| J114[c] | AGGTCGCCCGTGGTGGCCCA | NT | 121 |
| J116[c] | TGGTGGCCCATGGTCACCAT | NT | 131 |
| J118[c] | TGGTCACCATAGGTCACCCT | NT | 141 |
| J120[c] | AGGTCACCCTTGGCAACCAA | NT | 151 |
| hAAVS1[c] | GGGGCCACTAGGGACAGGAT | off-target | n/a |
| J206 | TAGTAGCCGAACACGTCCTC | off-target | n/a |
| J306 | TTGTGTCCAGAACGCTCCGT | NT | 81 |
| J306+1 | TGTGTCCAGAACGCTCCGTA | NT | 82 |
| J306+2 | GTGTCCAGAACGCTCCGTAG | NT | 83 |
| J306+3 | TGTCCAGAACGCTCCGTAGG | NT | 84 |
| J306+4 | GTCCAGAACGCTCCGTAGGG | NT | 85 |
| J306+5 | TCCAGAACGCTCCGTAGGGG | NT | 86 |

| | | | |
|---|---|---|---|
| M1 | AGCAGAAGTGTCAGCAGTGT | NT | 81 (reporter A), 86 (reporter B) |
| M2 | CGACGAGCAGAAGTGTCAGC | NT | 76 (reporter A), 81 (reporter B) |
| aroKB_A1 | GGGCAATTATTTCGTCATGA | T | 151 |
| aroKB_A2 | AGATGAACGACGCGAGTTAG | T | 122 |
| aroKB_A3 | TTTTACGGCTGTTTACTCAC | NT | 92 |
| aroKB_A4 | TGAGTAAACAGCCGTAAAAG | T | 71 |
| cysK_C1 | CCACCCCTGTTTCACACAAA | NT | 93 |
| cysK_C2 | AAACCGTTTGTGTGAAACAG | T | 79 |
| cysK_C3 | GACATGCAAGATGGAATAAG | NT | 61 |
| cysK_C4 | ATGACATGCAAGATGGAATA | NT | 59 |
| cysK_C5 | GGAAATAATGACATGCAAGA | NT | 52 |
| ldhA_L1 | CAGTAATAACAGCGCGAGAA | T | 157 |
| ldhA_L2 | GGATATTAACTACCCATGCT | NT | 140 |
| ldhA_L3 | GCTTTATATTTACCCAGCAT | T | 135 |
| ldhA_L4 | GCTTAATTTTTCGCTAAATC | NT | 119 |
| ldhA_L5 | GAAAAATTAAGCATTCAATA | T | 91 |
| ldhA_L6 | AGCATTCAATACGGGTATTG | T | 82 |
| ldhA_L7 | AGGCGCAACCTTCAACTGAA | NT | 67 |
| ldhA_L8 | ATGTTTAACCGTTCAGTTGA | T | 58 |
| yajG_Y1 | TTGACGAAATAATCGCCCCT | NT | 81 |
| yajG_Y2 | CATCAGTGTTTCTTTTACCA | T | 80 |
| yajG_Y3 | AAATAATCGCCCCTGGTAAA | NT | 87 |
| poxB_P1 | CCCGATGAAAGGAATATCAT | NT | 91 |
| poxB_P2 | GGTTAAATAGCCCGATGAAA | NT | 81 |
| uxuR_U1 | TGATTGACCAGTAAGTCTGT | NT | 81 |
| uxuR_U2 | GATTACCCTACAGACTTACT | T | 70 |
| ppiD_D1 | ACTAAGCGTTGTCCCCAGTG | T | 80 |
| ppiD_D2 | GTCCCCAGTGGGGATGTGAC | T | 70 |
| ansB_B1 | AGATCTACAAAGTTAGAGGC | NT | 91 |
| ansB_B2 | TATATTTTGGAGATCTACAA | NT | 81 |
| araE_E1 | TGCGACATGTCGTTATGTGA | NT | 91 |

| | | | |
|---|---|---|---|
| araE_E2 | ATTAAATTGCTGCGACATGT | NT | 81 |

[a] Template strand (T) or non-template strand (NT).

[b] Distance to TSS is the distance from the 3' end (PAM proximal) of the guide target site to the transcription start site. For synthetic promoters driven by BBa_J23117 or BBa_J23119 (http://parts.igem.org), the TSS is immediately downstream of the BBa sequence (see complete maps below).

[c] The J101-J121 sites were the same target sites used to test the positioning dependence of CRISPRa at a 10 bp resolution on the J1-J23117 promoter[1].

**Supplementary Table 4.** Select *E. coli* Expression Plasmids[a]

| **Plasmid** | **Marker** | **origin** | **Promoter** | **Gene** | **Terminator** |
|---|---|---|---|---|---|
| pCD442 | *CmR* | *p15A* | *1) Sp.pCas9*<br>*2) BBa_J23107* | 1) dCas9<br>2) MCP-(5aa)-SoxS (R93A/S101A) | *1) BBa_B0015*<br>*2) BBa_B1002* |
| pCK005.1-21[b] | *CmR* | *p15A* | *1) Sp.pCas9*<br>*2) BBa_J23107*<br>*3) BBa_J23119* | 1) dCas9<br>2) MCP-(5aa)-SoxS (R93A/S101A)<br>3) 1x MS2 scRNA.b2 (J101-121 targets) | *1) BBa_B0015*<br>*2) BBa_B1002*<br>*3) TrrnB* |
| pCD564 | *CmR* | *p15A* | *1) Sp.pCas9*<br>*2) BBa_J23107* | 1) dxCas9(3.7)<br>2) MCP-(5aa)-SoxS (R93A/S101A) | *1) BBa_B0015*<br>*2) BBa_B1002* |
| pCD565 | *CmR* | *p15A* | *1) Sp.pCas9*<br>*2) BBa_J23107*<br>*3) BBa_J23119* | 1) dxCas9(3.7)<br>2) MCP-(5aa)-SoxS (R93A/S101A)<br>3) 1x MS2 scRNA.b2 (J306 target) | *1) BBa_B0015*<br>*2) BBa_B1002*<br>*3) TrrnB* |
| pCD580.-1~-5 | *AmpR* | *pSC101 *** | *J3_BBa_J23117 (with 1-5 bp deleted from the J1 region)* | mRFP1 | *BBa_B0015* |
| pCD581 | *CmR* | *p15A* | *1) Sp.pCas9*<br>*2) BBa_J23107*<br>*3) BBa_J23119* | 1) dCas9<br>2) MCP-(5aa)-SoxS (R93A/S101A)<br>3) 1x MS2 scRNA.b2 (J306 target) | *1) BBa_B0015*<br>*2) BBa_B1002*<br>*3) TrrnB* |
| pJF076Sa[c] | *AmpR* | *pSC101 *** | *J3_BBa_J23117* | mRFP1 | *BBa_B0015* |
| pJF143-J3[d] | *AmpR* | *pSC101 *** | *J3_BBa_J23117* | mRFP1 | *BBa_B0015* |

| | | | | | |
|---|---|---|---|---|---|
| pJF155.1-12[c] | *AmpR* | *pSC101 *** | *J1_BBa_J23117 (with 1-12 bp inserted upstream of -35)* | mRFP1 | *BBa_B0015* |
| pJF161.1-12[d] | *AmpR* | *pSC101 *** | *J3_BBa_J23117 (with 1-12 bp inserted upstream of -35)* | mRFP1 | *BBa_B0015* |

[a] BBa sequences are from the Repository of Standard Biological Parts (http://parts.igem.org). dCas9 is the catalytically inactive form of *S. pyogenes* Cas9. Sp.pCas9 is the endogenous Cas9 promoter from *S. pyogenes*.

[b] pCK005.1-21 indicates a set of plasmids (pCK005.1, pCK005.2…) where the final number corresponds to guide RNA target sites (J101, J102…, Supplementary Table 3) used for the J1-117-mRFP1 reporter.

[c] Originally described in previous work[1]. Modified versions of this plasmid are available where BBa_J23117 is replaced with minimal promoters regulated by alternative sigma factors (Figure 3B).

[d] Modified version of this plasmid are available where: BBa_J23117 is replaced with Anderson promoters of different strength (Figure 3A) and different PAM sites at the -81 J306 site (Figure 5A).

**Supplementary Table 5.** Primer Sequences for RT-qPCR

| Primer | Sequence | Reference |
|---|---|---|
| 16S_f | AAAGTTAATACCTTTGCTCATTGACGTT | [1] |
| 16S_r | GACTACCAGGGTATCTAATCCTGTTT | [1] |
| aroK_f | TCTGGTTGGGCCTATGGGTG | This study |
| aroK_r | TACGAATGGTCACGTCGGCA | This study |
| cysK_f | TGCTGAAACCAGGCGTTGAA | This study |
| cysK_r | TCCCAACGCCAGCAATAAATACA | This study |
| ldhA_f | TGGCTGCGAAGCGGTATGTA | This study |
| ldhA_r | GAACGCCAGCAGACGCATAC | This study |
| yajG_fw | AAGTCACCCGCGATAAT | This study |
| yajG_rev | CTTTGGTCGCGATGTTG | This study |
| poxB_fw | GGTCTTAGTGACAGTCTTAATC | This study |
| poxB_r | GGAATATGAGCGGCAATC | This study |

**Supplementary methods**

Computational analysis of PAM site availability

The intergenic sequences from *E. coli* were obtained from the RegulonDB database[9]. To obtain the intergenic sequences upstream of promoters, the intergenic sequences between convergent genes and between intra-operon coding sequences were removed. 5' untranslated regions (UTRs) were removed from sequences using transcription start site (TSS) information from Kim et al.[10] and RegulonDB. Intergenic sequences upstream of genes with no known TSS or where no intergenic sequence remained after removing the UTRs were discarded, yielding 1504 transcriptional units for further analysis. The number of PAM sites was calculated by counting the number of PAM sites on both strands (Supplementary Figure 8D). The PAM density over a 10 bp window was calculated by dividing the number of PAMs found at each intergenic sequence by the length of the intergenic sequence and multiplying by 10. Analyses were performed using Python 2.7.

Computational selection of candidate endogenous genes for CRISPRa

PAM sites found in *E. coli* intergenic sequences upstream of promoters were assigned a "sequence score" and a "distance score". The sequence score indicates the relative activity of a given PAM sequence compared to an AGG PAM, and was calculated using data from Figure 5A with dxCas9(3.7) (e.g. AGG = 1, CGT = 0.24). All NGG sequences were assigned a "sequence score" of 1. The distance score indicated the relative activity of a given PAM site position compared to a PAM site found at -81 on the non-template strand or -70 on the template from the TSS. Distance scores for the non-template strand were calculated using data from Figure 4A (e.g. -81 = 1, -91 = 0.62). For the template strand, the same scores were used, but each score was shifted upstream by 11 bases to match the position of the site of maximum activation (e.g. -70 = 1, -80 = 0.62). Each PAM site was assigned a final calculated score as "sequence score" x "position score". Promoters were then ranked by sorting for higher PAM scores at the peaks of activation (sum of the scores for the PAMs at -70, -80, -81, -91). This analysis was performed using Python 2.7. Candidates were then manually selected from among the top scoring promoters by the following criteria: (1) two or more candidate PAM sites, (2) regulation by $\sigma^{70}$, and (3) relatively weak basal expression level (<10% of the level of the maximally expressed *E. coli* gene according to data from the *E. coli* promoter collection[8]. Using these criteria, we chose the following six candidates for further characterization: *yajG*, *uxuR*, *ansB*, *poxB*, *araE*, and *ppiD*. Each of these genes is represented in the *E. coli* promoter collection (Dharmacon), a commercially available library of promoter-GFPmut2 fusions[8].

**Sequence of the double mutant activation domain: MCP-(5aa)-SoxS(R93A/S101A)**

> MCP-(5aa)-SoxS(R93A/S101A) (optimized for minimal endogenous activity, Figure 1 and subsequent)

MCP$_{\Delta FG,\ V29I}$., 5 aa linker, SoxS(R93A/S101A) (underlined are the alanine point mutations)

MGPASNFTQFVLVDNGGTGDVTVAPSNFANGIAEWISSNSRSQAYKVTCSVRQSSAQNRKYTIKVEVPKG
AWRSYLNMELTIPIFATNSDCELIVKAMQGLLKDGNPIPSAIAANSGIYGGGGSMSHQKIIQDLIAWIDE
HIDQPLNIDVVAKKSGYSKWYLQRMFRTVTHQTLGDYIRQRRLLLAAVELRTTERPIFDIAMDLGYVSQQ
TFSRVFARQFDRTPADYRHRL

**Reporter genes**

The integrated sfGFP reporter, the *zwfp-lacZ* and *fumCp-lacZ* reporters used for testing the CRISPRa and endogenous activities of mutant SoxS (Figure 1) and the J1-J23117-mRFP1 reporter (Figure 3, Supplementary Figure 8B) for testing the distance dependence property were described in previous study[1].

J3-J23117_mRFP1 reporter (Figure 3 & S2 and subsequent)

The J3 upstream sequence contains a PAM site allowing guide RNAs to target at -81 bp to the TSS on the non-template strand, which is the same distance where maximum CRISPRa activity was observed on the J1 upstream region described previously[1].

J3 upstream region, BBa_J23117 promoter, Bujard RBS, mRFP1, BBa_B0015 terminator. The J306 target site is underlined.

AGCATTTGCGATCATTCACGCAGCGCTTATTCAGTTGCTCACTGCGATGTCATAATCATCGCTACGAGCT
GTGAAAGATGCATAAAGCTCGTACGACGCGTTCGCTCGTCTCCTCACTTCTCCTACGGAGCGTTCTGGAC
ACAACGTCGTCTTGAAGTTGCGATTATAGATTGACAGCTAGCTCAGTCCTAGGGATTGTGCTAGCGAATT
CATTAAAGAGGAGAAAGGTACCATGGCGAGTAGCGAAGACGTTATCAAAGAGTTCATGCGTTTCAAAGTT
CGTATGGAAGGTTCCGTTAACGGTCACGAGTTCGAAATCGAAGGTGAAGGTGAAGGTCGTCCGTACGAAG
GTACCCAGACCGCTAAACTGAAAGTTACCAAAGGTGGTCCGCTGCCGTTCGCTTGGGACATCCTGTCCCC
GCAGTTCCAGTACGGTTCCAAAGCTTACGTTAAACACCCGGCTGACATCCCGGACTACCTGAAACTGTCC
TTCCCGGAAGGTTTCAAATGGGAACGTGTTATGAACTTCGAAGACGGTGGTGTTGTTACCGTTACCCAGG
ACTCCTCCCTGCAAGACGGTGAGTTCATCTACAAAGTTAAACTGCGTGGTACCAACTTCCCGTCCGACGG
TCCGGTTATGCAGAAAAAAACCATGGGTTGGGAAGCTTCCACCGAACGTATGTACCCGGAAGACGGTGCT

CTGAAAGGTGAAATCAAAATGCGTCTGAAACTGAAAGACGGTGGTCACTACGACGCTGAAGTTAAAACCA
CCTACATGGCTAAAAAACCGGTTCAGCTGCCGGGTGCTTACAAAACCGACATCAAACTGGACATCACCTC
CCACAACGAAGACTACACCATCGTTGAACAGTACGAACGTGCTGAAGGTCGTCACTCCACCGGTGCTTAA
GGATCCAAACTCGAGTAAGGATCTCCAGGCATCAAATAAAACGAAAGGCTCAGTCGAAAGACTGGGCCTT
TCGTTTTATCTGTTGTTTGTCGGTGAACGCTCTCTACTAGAGTCACACTGGCTCACCTTCGGGTGGGCCT
TTCTGCGTTTATA

J3(N26)-J23117-sfGFP reporter library (Figure 3)

J3 upstream region (without 26 bases at 3'), N26, BBa_J23117 promoter, Bujard RBS, sfGFP, BBa_B0015 terminator. The J306 target site is underlined.

AGCATTTGCGATCATTCACGCAGCGCTTATTCAGTTGCTCACTGCGATGTCATAATCATCGCTACGAGCT
GTGAAAGATGCATAAAGCTCGTACGACGCGTTCGCTCGTCTCCTCACTTCTCCTACGGAGCGTTCTGGAC
ACAANNNNNNNNNNNNNNNNNNNNNNNNNNTTGACAGCTAGCTCAGTCCTAGGGATTGTGCTAGCGAATT
CATTAAAGAGGAGAAAGGTACCATGAGCAAAGGAGAAGAACTTTTCACTGGAGTTGTCCCAATTCTTGTT
GAATTAGATGGTGATGTTAATGGGCACAAATTTTCTGTCCGTGGAGAGGGTGAAGGTGATGCTACAAACG
GAAAACTCACCCTTAAATTTATTTGCACTACTGGAAAACTACCTGTTCCGTGGCCAACACTTGTCACTAC
TCTGACCTATGGTGTTCAATGCTTTTCCCGTTATCCGGATCACATGAAACGGCATGACTTTTTCAAGAGT
GCCATGCCCGAAGGTTATGTACAGGAACGCACTATATCTTTCAAAGATGACGGGACCTACAAGACGCGTG
CTGAAGTCAAGTTTGAAGGTGATACCCTTGTTAATCGTATCGAGTTAAAGGGTATTGATTTTAAAGAAGA
TGGAAACATTCTTGGACACAAACTCGAGTACAACTTTAACTCACACAATGTATACATCACGGCAGACAAA
CAAAAGAATGGAATCAAAGCTAACTTCAAAATTCGCCACAACGTTGAAGATGGTTCCGTTCAACTAGCAG
ACCATTATCAACAAAATACTCCAATTGGCGATGGCCCTGTCCTTTTACCAGACAACCATTACCTGTCGAC
ACAATCTGTCCTTTCGAAAGATCCCAACGAAAAGCGTGACCACATGGTCCTTCTTGAGTTTGTAACTGCT
GCTGGGATTACACATGGCATGGATGAGCTCTACAAATAAGGATCCAAACTCGAGTAAGGATCTCCAGGCA
TCAAATAAAACGAAAGGCTCAGTCGAAAGACTGGGCCTTTCGTTTTATCTGTTGTTTGTCGGTGAACGCT
CTCTACTAGAGTCACACTGGCTCACCTTCGGGTGGGCCTTTCTGCGTTTATA

**Modified synthetic promoters**

Promoters with different basal expression levels (Figure 3)

Annotations:

J3 upstream region, variable promoter (BBa_J231xx), Bujard RBS, Start codon of mRFP. The J106 target site is underlined

BBa_J23105

AGCATTTGCGATCATTCACGCAGCGCTTATTCAGTTGCTCACTGCGATGTCATAATCATCGCTACGAGCT
GTGAAAGATGCATAAAGCTCGTACGACGCGTTCGCTCGTCTCCTCACTTCTCCTACGGAGCGTTCTGGAC
ACAACGTCGTCTTGAAGTTGCGATTATAGAtttacggctagctcagtcctaggtactatgctagcGAATT
CATTAAAGAGGAGAAAGGTACCATG

BBa_J23106

AGCATTTGCGATCATTCACGCAGCGCTTATTCAGTTGCTCACTGCGATGTCATAATCATCGCTACGAGCT
GTGAAAGATGCATAAAGCTCGTACGACGCGTTCGCTCGTCTCCTCACTTCTCCTACGGAGCGTTCTGGAC
ACAACGTCGTCTTGAAGTTGCGATTATAGAtttacggctagctcagtcctaggtatagtgctagcGAATT
CATTAAAGAGGAGAAAGGTACCATG

BBa_J23107

AGCATTTGCGATCATTCACGCAGCGCTTATTCAGTTGCTCACTGCGATGTCATAATCATCGCTACGAGCT
GTGAAAGATGCATAAAGCTCGTACGACGCGTTCGCTCGTCTCCTCACTTCTCCTACGGAGCGTTCTGGAC
ACAACGTCGTCTTGAAGTTGCGATTATAGAtttacggctagctcagccctaggtattatgctagcGAATT
CATTAAAGAGGAGAAAGGTACCATG

BBa_J23108

AGCATTTGCGATCATTCACGCAGCGCTTATTCAGTTGCTCACTGCGATGTCATAATCATCGCTACGAGCT
GTGAAAGATGCATAAAGCTCGTACGACGCGTTCGCTCGTCTCCTCACTTCTCCTACGGAGCGTTCTGGAC
ACAACGTCGTCTTGAAGTTGCGATTATAGActgacagctagctcagtcctaggtataatgctagcGAATT
CATTAAAGAGGAGAAAGGTACCATG

BBa_J23109

AGCATTTGCGATCATTCACGCAGCGCTTATTCAGTTGCTCACTGCGATGTCATAATCATCGCTACGAGCT
GTGAAAGATGCATAAAGCTCGTACGACGCGTTCGCTCGTCTCCTCACTTCTCCTACGGAGCGTTCTGGAC
ACAACGTCGTCTTGAAGTTGCGATTATAGAtttacagctagctcagtcctagggactgtgctagcGAATT
CATTAAAGAGGAGAAAGGTACCATG

BBa_J23110

AGCATTTGCGATCATTCACGCAGCGCTTATTCAGTTGCTCACTGCGATGTCATAATCATCGCTACGAGCT
GTGAAAGATGCATAAAGCTCGTACGACGCGTTCGCTCGTCTCCTCACTTCTCCTACGGAGCGTTCTGGAC

ACAACGTCGTCTTGAAGTTGCGATTATAGAtttacggctagctcagtcctaggtacaatgctagcGAATT
CATTAAAGAGGAGAAAGGTACCATG

BBa_J23111
AGCATTTGCGATCATTCACGCAGCGCTTATTCAGTTGCTCACTGCGATGTCATAATCATCGCTACGAGCT
GTGAAAGATGCATAAAGCTCGTACGACGCGTTCGCTCGTCTCCTCACTTCTCCTACGGAGCGTTCTGGAC
ACAACGTCGTCTTGAAGTTGCGATTATAGAttgacggctagctcagtcctaggtatagtgctagcGAATT
CATTAAAGAGGAGAAAGGTACCATG

BBa_J23112
AGCATTTGCGATCATTCACGCAGCGCTTATTCAGTTGCTCACTGCGATGTCATAATCATCGCTACGAGCT
GTGAAAGATGCATAAAGCTCGTACGACGCGTTCGCTCGTCTCCTCACTTCTCCTACGGAGCGTTCTGGAC
ACAACGTCGTCTTGAAGTTGCGATTATAGActgatagctagctcagtcctagggattatgctagcGAATT
CATTAAAGAGGAGAAAGGTACCATG

BBa_J23113
AGCATTTGCGATCATTCACGCAGCGCTTATTCAGTTGCTCACTGCGATGTCATAATCATCGCTACGAGCT
GTGAAAGATGCATAAAGCTCGTACGACGCGTTCGCTCGTCTCCTCACTTCTCCTACGGAGCGTTCTGGAC
ACAACGTCGTCTTGAAGTTGCGATTATAGActgatggctagctcagtcctagggattatgctagcGAATT
CATTAAAGAGGAGAAAGGTACCATG

BBa_J23114
AGCATTTGCGATCATTCACGCAGCGCTTATTCAGTTGCTCACTGCGATGTCATAATCATCGCTACGAGCT
GTGAAAGATGCATAAAGCTCGTACGACGCGTTCGCTCGTCTCCTCACTTCTCCTACGGAGCGTTCTGGAC
ACAACGTCGTCTTGAAGTTGCGATTATAGAtttatggctagctcagtcctaggtacaatgctagcGAATT
CATTAAAGAGGAGAAAGGTACCATG

BBa_J23115
AGCATTTGCGATCATTCACGCAGCGCTTATTCAGTTGCTCACTGCGATGTCATAATCATCGCTACGAGCT
GTGAAAGATGCATAAAGCTCGTACGACGCGTTCGCTCGTCTCCTCACTTCTCCTACGGAGCGTTCTGGAC
ACAACGTCGTCTTGAAGTTGCGATTATAGAtttatagctagctcagcccttggtacaatgctagcGAATT
CATTAAAGAGGAGAAAGGTACCATG

BBa_J23118

AGCATTTGCGATCATTCACGCAGCGCTTATTCAGTTGCTCACTGCGATGTCATAATCATCGCTACGAGCT
GTGAAAGATGCATAAAGCTCGTACGACGCGTTCGCTCGTCTCCTCACTTCTCCTACGGAGCGTTCTGGAC
ACAACGTCGTCTTGAAGTTGCGATTATAGAttgacggctagctcagtcctaggtattgtgctagcGAATT
CATTAAAGAGGAGAAAGGTACCATG

BBa_J23119

AGCATTTGCGATCATTCACGCAGCGCTTATTCAGTTGCTCACTGCGATGTCATAATCATCGCTACGAGCT
GTGAAAGATGCATAAAGCTCGTACGACGCGTTCGCTCGTCTCCTCACTTCTCCTACGGAGCGTTCTGGAC
ACAACGTCGTCTTGAAGTTGCGATTATAGAttgacagctagctcagtcctaggtataatgctagcGAATT
CATTAAAGAGGAGAAAGGTACCATG

Promoters regulated by alternative sigma factors (Figure 3)

Annotations:

J1 upstream region, variable promoter (color coded), Bujard RBS, Start codon of mRFP. The J106 target site is underlined

J1-*sodCp* promoter

GCCTACGGTATCCACCGGAGACCTATGGCAGCCTCCGGCCGCCATAGGACACCTTTGGTTGCCAAGGGTG
ACCTATGGTGACCATGGGCCACCACGGGCGACCTCAGGTATCCTGCGGTGTCCTGCGGTTACCAAAGGCG
TCCTTTGGGTTCCACCGGATACCTCCGGACGCCGTTcaaaaatgtgtcacTGGTTTACACTtattcagGG
AATTCATTAAAGAGGAGAAAGGTACCATG

J1-*glnAp2* promoter

GCCTACGGTATCCACCGGAGACCTATGGCAGCCTCCGGCCGCCATAGGACACCTTTGGTTGCCAAGGGTG
ACCTATGGTGACCATGGGCCACCACGGGCGACCTCAGGTATCCTGCGGTGTCCTGCGGTTACCAAAGGCG
TCCTTTGGGTTCCACCGGATACCTCCGGACTGGCACagatttCGCTTtatctttttTacggcgacGAATT
CATTAAAGAGGAGAAAGGTACCATG

J1-*rdgBp* promoter

GCCTACGGTATCCACCGGAGACCTATGGCAGCCTCCGGCCGCCATAGGACACCTTTGGTTGCCAAGGGTG
ACCTATGGTGACCATGGGCCACCACGGGCGACCTCAGGTATCCTGCGGTGTCCTGCGGTTACCAAAGGCG
TCCTTTGGGTTCCACCGGATACCTCCGGACTTGTAAaaggcgaacttggcCGCCACaaacaaattGAATT
CATTAAAGAGGAGAAAGGTACCATG

J1-*yieEp* promoter

GCCTACGGTATCCACCGGAGACCTATGGCAGCCTCCGGCCGCCATAGGACACCTTTGGTTGCCAAGGGTG
ACCTATGGTGACCATGGGCCACCACGGGCGACCTCAGGTATCCTGCGGTGTCCTGCGGTTACCAAAGGCG
TCCTTTGGGTTCCACCGGATACCTCCGGACGAACTTttagccgctttagtctgtcCATCAttccAGAATT
CATTAAAGAGGAGAAAGGTACCATG

J3(tetO)-J23117 mRFP1 reporter (Figure 3)

J3 upstream region (without 19 bases at 3'), tetO, BBa_J23117 promoter, Bujard RBS, ATG of mRFP1. The J306 target site is underlined.

AGCATTTGCGATCATTCACGCAGCGCTTATTCAGTTGCTCACTGCGATGTCATAATCATCGCTACGAGCT
GTGAAAGATGCATAAAGCTCGTACGACGCGTTCGCTCGTCTCCTCACTTCTCCTACGGAGCGTTCTGGAC
ACAACGTCGTCACTCTATCGTTGATAGAGTttgacagctagctcagtcctagggattgtgctagcGAATT
CATTAAAGAGGAGAAAGGTACCATG

J1-J23117 promoter with shifted bases (Figure 4, Supplementary Figure S3 and subsequent)

For promoters shifted by fewer than 12bp, bases were removed from the 5' end of the inserted sequence (starting with C).

J1 upstream region, inserted sequence (12 bases), BBa_J23117 promoter, Bujard RBS, ATG of mRFP1. The J106 target site is underlined.

GCCTACGGTATCCACCGGAGACCTATGGCAGCCTCCGGCCGCCATAGGACACCTTTGGTTGCCAAGGGTG
ACCTATGGTGACCATGGGCCACCACGGGCGACCTCAGGTATCCTGCGGTGTCCTGCGGTTACCAAAGGCG
TCCTTTGGGTTCCACCGGATACCTCCGGACCTAGATGCGCTCttgacagctagctcagtcctagggattg
tgctagcGAATTCATTAAAGAGGAGAAAGGTACCATG

J3-J23117 promoter with shifted bases (Supplementary Figure 3 and subsequent)

For promoters shifted by fewer than 12bp, bases were removed from the 5' end of the inserted sequence (starting with T). For promoters with negative shifts, no sequence was inserted between the J3 upstream region and BBa_J23117, and bases were removed from the 3' end of the J3 upstream region (starting with A).

J3 upstream region, inserted sequence (12 bases), BBa_J23117 promoter, Bujard RBS, ATG of mRFP1. The J306 target site is underlined.

AGCATTTGCGATCATTCACGCAGCGCTTATTCAGTTGCTCACTGCGATGTCATAATCATCGCTACGAGCT
GTGAAAGATGCATAAAGCTCGTACGACGCGTTCGCTCGTCTCCTCACTTCTCCTACGGAGCGTTCTGGAC
ACAACGTCGTCTTGAAGTTGCGATTATAGATGAGATGCGCTCttgacagctagctcagtcctagggattg
tgctagcGAATTCATTAAAGAGGAGAAAGGTACCATG

Modified J3-J23117 promoter with 6 adjacent PAMs around -81 bp to TSS (Supplementary Figure 3)

J3 upstream region, BBa_J23117 promoter, Bujard RBS, Start codon of mRFP. Region with additional PAM sites inserted. The J306 target site is underlined.

AGCATTTGCGATCATTCACGCAGCGCTTATTCAGTTGCTCACTGCGATGTCATAATCATCGCTACGAGCT
GTGAAAGATGCATAAAGCTCGTACGACGCGTTCGCTCGTCTCCTCACCCCCCCTACGGAGCGTTCTGGAC
ACAACGTCGTCTTGAAGTTGCGATTATAGAttgacagctagctcagtcctagggattgtgctagcGAATT
CATTAAAGAGGAGAAAGGTACCATG

Modified J3-J23117 promoter with non-NGG PAMs around -81 bp to TSS (Figure 5A)

J3 upstream region, BBa_J23117 promoter, Bujard RBS, Start codon of mRFP, region with modified PAMs. The J306 target site is underlined.

AGCATTTGCGATCATTCACGCAGCGCTTATTCAGTTGCTCACTGCGATGTCATAATCATCGCTACGAGCT
GTGAAAGATGCATAAAGCTCGTACGACGCGTTCGCTCGTCTCCTCACTTCTNNNACGGAGCGTTCTGGAC
ACAACGTCGTCTTGAAGTTGCGATTATAGAttgacagctagctcagtcctagggattgtgctagcGAATT
CATTAAAGAGGAGAAAGGTACCATG

Promoter to demonstrate dxCas9(3.7) versatility (Figure 5B)

J3 upstream region, BBa_J23117 promoter, Bujard RBS, Start codon of mRFP. The modified target site (M) is underlined. AGT PAM site.

AGCATTTGCGATCATTCACGCAGCGCTTATTCAGTTGCTCACTGCGATGTCATAATCATCGCTACGAGCT
GTGAAAGATGCATAAAGCTCGTACGACGCGTTCGCTCGTCTCCTCACTTCTCCTACACTGCTGACACTTC
TGCTCGTCGTCTTGAAGTTGCGATTATAGAttgacagctagctcagtcctagggattgtgctagcGAATT
CATTAAAGAGGAGAAAGGTACCATG

**Supplementary references**

1. Dong, C., Fontana, J., Patel, A., Carothers, J. M. & Zalatan, J. G. Synthetic CRISPR-Cas gene activators for transcriptional reprogramming in bacteria. *Nat Commun* **9,** 2489 (2018).

2. Arai, R., Ueda, H., Kitayama, A., Kamiya, N. & Nagamune, T. Design of the linkers which effectively separate domains of a bifunctional fusion protein. *Protein Eng Des Sel* **14,** 529–532 (2001).

3. Zalatan, J. G. *et al.* Engineering Complex Synthetic Transcriptional Programs with CRISPR RNA Scaffolds. *Cell* **160,** 339–350 (2015).

4. Konermann, S. *et al.* Genome-scale transcriptional activation by an engineered CRISPR-Cas9 complex. *Nature* **517,** 583–588 (2015).

5. Bikard, D. *et al.* Programmable repression and activation of bacterial gene expression using an engineered CRISPR-Cas system. *Nucleic Acids Res* **41,** 7429–7437 (2013).

6. Griffith, K. L. & Wolf, R. E. A Comprehensive Alanine Scanning Mutagenesis of the Escherichia coli Transcriptional Activator SoxS: Identifying Amino Acids Important for DNA Binding and Transcription Activation. *J Mol Biol* **322,** 237–257 (2002).

7. Hu, J. H. *et al.* Evolved Cas9 variants with broad PAM compatibility and high DNA specificity. *Nature* **556,** 57–63 (2018).

8. Zaslaver, A. *et al.* A comprehensive library of fluorescent transcriptional reporters for Escherichia coli. *Nature Methods* **3,** 623–628 (2006).

9. Gama-Castro, S. *et al.* RegulonDB version 9.0: high-level integration of gene regulation, coexpression, motif clustering and beyond. *Nucleic Acids Res* **44,** D13–D143 (2016).

10. Kim, D. *et al.* Comparative Analysis of Regulatory Elements between Escherichia coli and Klebsiella pneumoniae by Genome-Wide Transcription Start Site Profiling. *PLoS Genetics* **8,** e1002867-15
